# Effect of nanocarbon molecules on the *Arabidopsis thaliana* transcriptome

**DOI:** 10.1101/2020.05.22.110171

**Authors:** Norihito Nakamichi, Ayato Sato, Yusuke Aihara, Toshinori Kinoshita, Yasutomo Segawa, Kazuma Amaike, Kenichiro Itami

## Abstract

Nanocarbons, such as fullerenes, carbon nanotubes, and graphene have attracted a great deal of attention as next-generation materials because of their unprecedented structures and unique physicochemical properties; however, almost all nanocarbons reported previously were used as mixtures. Thus, there are still many unsolved issues about their biological functions at the molecular level. Our synthetic campaign in the last decade has synthesized structurally uniform and atomically precise nanocarbons, leading to the preparation of a library consisting of eighty structurally diverse nanocarbon molecules. This resource motivated us to explore the as yet uncovered biological functions of these nanocarbon molecules in organisms. Recently, nanotubes were used to deliver genes to plants; however, the effects of the molecules on plants are not well known. To monitor the effects of nanocarbon molecules on plants, we analyzed the transcriptome of *Arabidopsis thaliana* seedlings treated with [9]cycloparaphenylene (CPP), decaborylated warped nanographene (WNG), and dimethoxyhexabenzotetracene (HBT). Clustering analysis indicated few effects of nanocarbon molecules on the transcriptome, perhaps suggesting a low toxicity of nanocarbon molecules on plants. We found that *AT1G05880* (*ARIADNE 12*) gene, categorized into ‘response to hypoxia’ genes, was up-regulated by nanocarbon molecules, suggesting that this gene is usable as maker for treatment of nanocarbon molecules.

## 1. Introduction

Carbon materials, as represented by fullerenes, carbon nanotubes, graphene, carbon nanobelts, and graphene nanoribbons, have attracted much attention as next-generation materials and have stimulated fundamental research in the fields of electronics, energy, and medical applications[1-3]. The use of nanocarbon molecules to analyze and control biological systems is also growing[4]. For example, functionalized fullerenes were used for gene delivery in mice[5]. Recent studies in which some nanocarbon molecules, carbon nanotubes (CNTs) or graphene oxide, accelerated gene transfection suggested that nanocarbons, although their biomedical properties occupied in the chemical space could be far off from the one of druggable molecules, could also possess some specific biological target molecules[6]. In addition, their unique structures and physicochemical properties have also been proposed to control not only target biomolecules but also dynamics in mesoscopic areas such as the interaction of biomolecules e.g., protein-protein interaction, membrane trafficking, and cell-cell communications. Indeed, graphene has positive effects such as inducing cell growth and causing a shielding effect by binding to plasma membranes in mammalian cells but with some concurrent damage to plasma membranes[7]. Almost all nanocarbon molecules reported previously have been delivered as a mixture; thus, there remain many unsolved issues about their precise biological functions at the molecular level. Our synthetic campaign in the past decade has succeeded in the syntheses of ‘structurally uniform’ and ‘atomically precise’ nanocarbons[2,8-10], giving us an opportunity to explore precisely their functions in organisms. For example, cell death of HeLa cells was induced using a uniform and water-soluble warped nanographene irradiated with a 489 nm laser, thereby providing new methodology to control cell activity[11].

Although much attention has been given to the use of nanocarbons in animal biology or medical sciences, nanocarbon applications in the plant science field are also emerging. For example, functionalized nanotubes have been used for chloroplast-selective gene delivery[12]. Functionalized high-aspect-ratio nanotubes and nanoparticles provide an efficient means of gene delivery to some plants[13]. Land plants cannot escape from an unfavorable environment by moving as animals can. Plants must respond to the environment where they germinate by coordinating their metabolic and physiological activities with environmental fluctuations. The ability to regulate genome-wide gene expression in response to specific environmental changes is generally crucial for plant adaptation. Genome-wide gene regulation is already triggered before plants initiate their physiological responses to environmental changes. It is possible to regard alternation of gene expression as ‘marker’ that represents one of plant responses to environmental changes.

To understand the physiological responses of plants to nanocarbon molecules, we analyzed the transcriptome of *Arabidopsis thaliana* (Arabidopsis) seedlings that were treated with three representative nanocarbon molecules. Clustering analysis indicated that the overall effect of these nanocarbon molecules was weak on the transcriptome, perhaps suggesting low toxicity of the nanocarbon molecules to Arabidopsis. We found that accumulation of *AT1G05880 (ARIADNE 12*) mRNA was increased by treatment of two nanocarbon molecules.

## 2. Results

### 2.1. RNAseq sample for plant treated with nanocarbon molecules

To gain insight into nanocarbons-effects on plants, we chose three nanocarbon molecules (CPP[1], WNG[14], and HBT[15]) from our ITbM nanocarbon library, as model molecules (Figure 1A). *Arabidopsis thaliana* (Arabidopsis) was chosen because the Arabidopsis genome information and many archive transcriptome data are available. In addition, we have established the method to treat Arabidopsis seedlings with small molecules (compounds) and analyze transcriptome[16]. A transcriptome is often already altered before organisms start to the physiological response to the treatment. Thus, transcriptome analyses of plants treated with small molecules provided insight into action mechanism of the molecules[17-19]. Thus, we conducted a genome-wide gene expression analysis to understand plant responses to nanocarbon molecules. Young seedling is suitable for treatment and presumably sensitive to any treatments, because epidermal of cotyledons are structurally soft than that of true leaves. Due to the limitation of the quantity of nanocarbon molecules as material, we designed the RNAseq experiment as below (Figure 1B). Since gathering seedlings into one sample presumably causes averaging of the transcriptome of the seedlings, which could improve data integrity if biological replicates are not available. Due to lacking information on treatment duration, we planned to sample seedlings in three time points. Nanocarbon molecules were treated to twenty-four seedlings and respective eight seedlings were harvested as one sample at 3 h, 9 h, or 24 h after the treatment and transcriptome change were analyzed (Figure 1B-1E). Finally, we validated RNAseq data by RT-qPCR using different biological samples.

**Figure 1.**
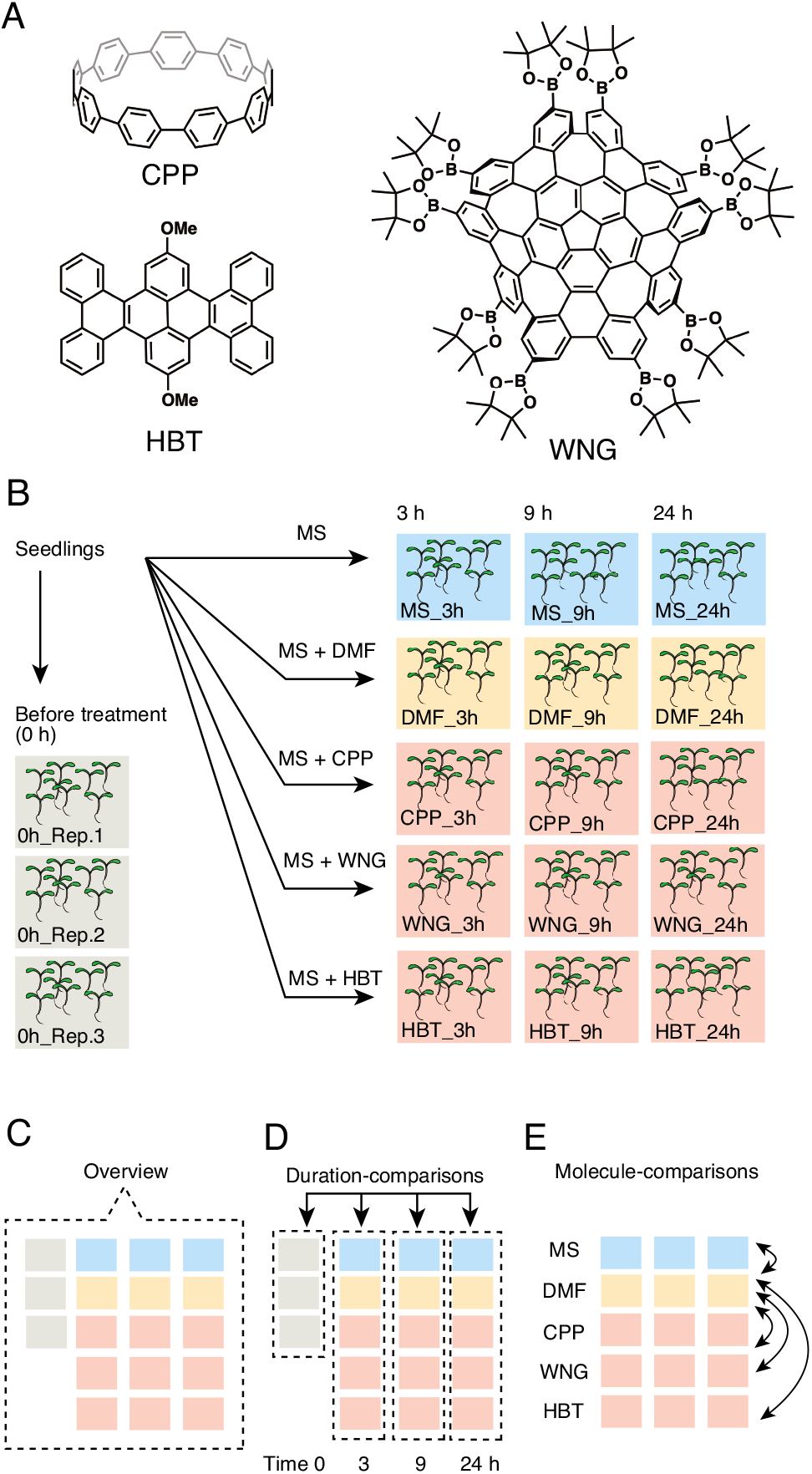
Design of transcriptome analyses for Arabidopsis treated with nanocarbon molecules. (**A**) Structures of nanocarbon molecules. CPP, WNG, and HBT. (**B**) Scheme of sampling for transcriptome analysis. Seedlings were treated with MS liquid, MS liquid containing DMF, CPP, WNG, or HBT. Seedlings were sampled at 3, 9, and 24 h after treatment. (**C**) The overview shows the result of a clustering analysis using whole transcriptome data. (**D**) Detection of treatment duration-dependent gene expression changes. (**E**) Detection of nanocarbon-dependent gene expression changes.

Arabidopsis seedlings grown under constant light conditions were treated with 50 μM of CPP, WNG, HBT, the solvent DMF (N,N-dimethylformamide) solved in MS (Murashige-Skoog) media, or without DMF (MS) (Figure 1B). Whole organs (cotyledons, hypocotyl, and root) was exposed to the treatment. It was demonstrated that 1-50 μM of bioactive small molecules change gene expression in Arabidopsis seedlings[17,18,20]. Thus we assumed that nanocarbon molecules may affect gene expression if these molecules have any biological activities. Three, 9, and 24 h after the treatment, eight seedlings were gathered as a sample in a test tube, and frozen by liquid nitrogen. We obtained at least 10,000,000 reads per sample (Supplemental Table 1). Sequence quality values Q20 and Q30 were over 90, and mapping rates to the TAIR 10 gene model (http://www.arabidopsis.org/) were over 95% (Supplemental Table1). We concluded that read quantity and quality of RNAseq data were enough to understand transcriptome change by the treatment of nanocarbon molecules. The expression level of each gene was normalized counts per million (CPM) (Supplemental Dataset1).

### 2.2. Analyses of RNAseq data

Although individual seedlings are considered to have a circadian rhythm or other developmental timers even under constant light conditions, our strategy was to pool eight seedlings into one sample, thereby attenuating time information by desynchronizing the clock and timer of individual seedlings. To test this hypothesis, we analyzed the expression of high-amplitude circadian clock genes *CIRCADIAN CLOCK-ASSOCIATED 1* (*CCA1*)[21], *TIMING OF CAB EXPRESSION 1*(*TOC1*)[22,23], and *GIGANTEA* (*GI*)[24] in our RNAseq data. Expression of these three genes was relatively constant throughout the time course of the experiment 0, 3, 9, and 24 h (the ratio of maximal expression/ minimal expression was smaller than 3) (Supplementary Figure 1). Constant expression of these genes was observed in plants treated with nanocarbon molecules (Supplementary Figure 1). These results suggested that the circadian clocks of seedlings comprising a sample were desynchronized, and that clock gene expression is averaged; thus, circadian time can be almost ignored in our samples.

To obtain an overview of genome-wide gene expression, all RNAseq data were examined by clustering analysis. We found that the ‘duration of treatment’ was a major factor for distinguishing whole transcriptome profiles (Figure 2). To identify genes whose expression was altered in a duration-dependent manner, we compared the transcriptomes of samples from treatment times of 0, 3, 9, and 24 h (Figure 3A). The number of duration-dependent mis-expressed genes are shown (EdgeR, False discovery rate FDR controlled q < 10^−10^).

**Figure 2.**
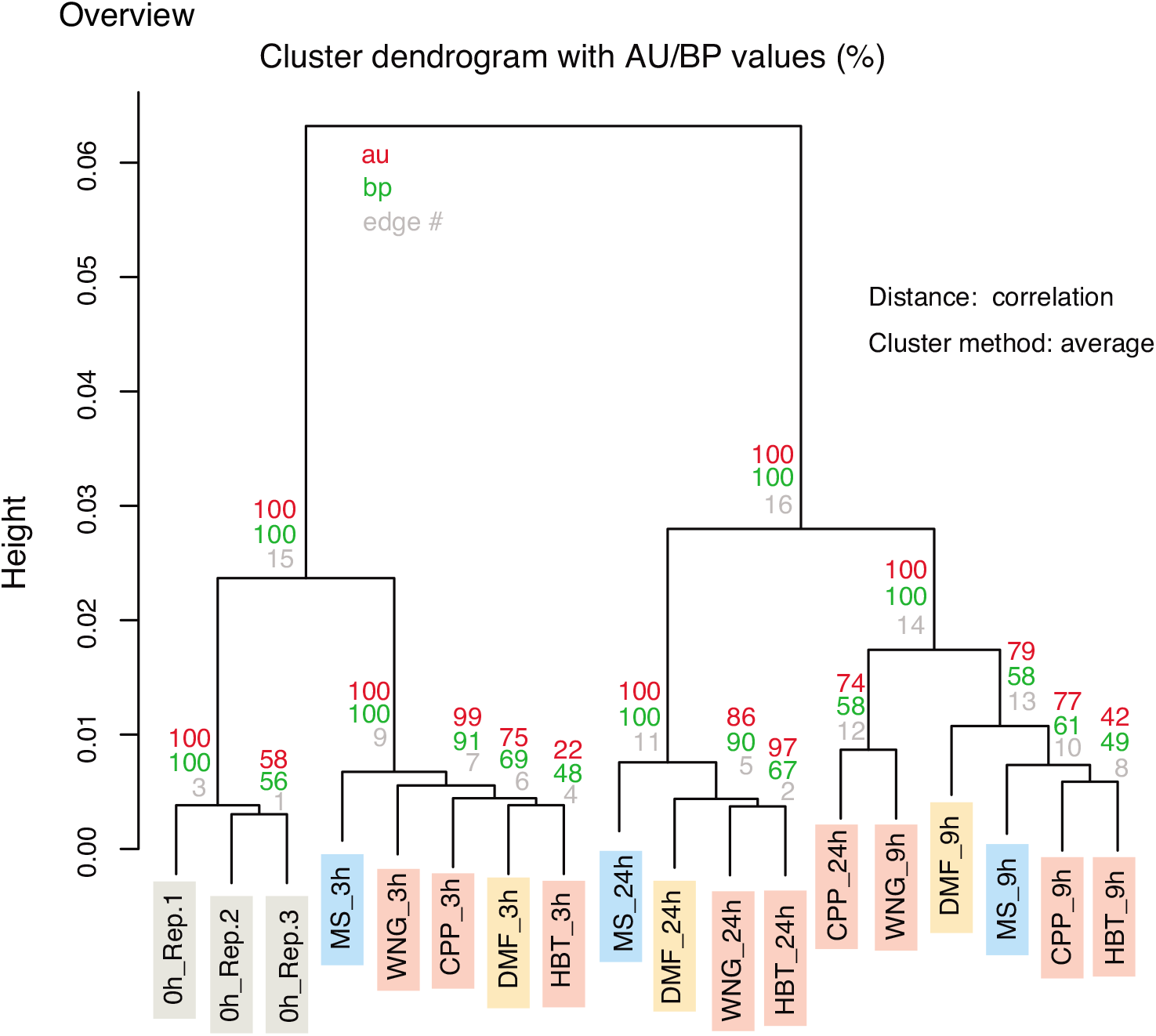
Clustering analysis of whole transcriptome data. au and bp mean approximately unbiased *p*-value and bootstrap probability value, respectively.

**Figure 3.**
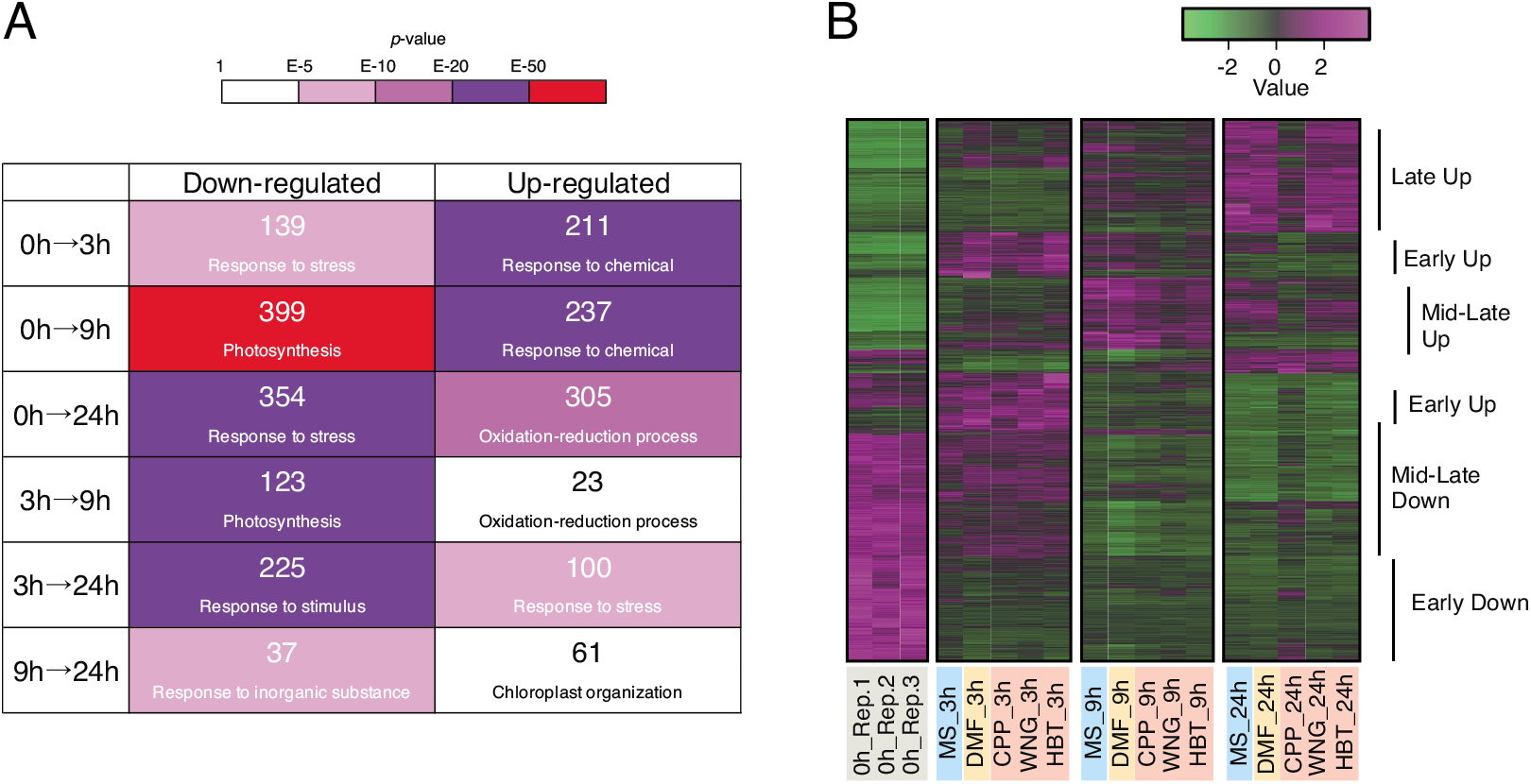
Treatment duration-dependent alteration of gene expression. (**A)** Number of genes whose expression was statistically changed among different sampling times (EdgeR, False discovery rate FDR controlled *q* < 10^−10^) is shown as the upper value in each box. The best hit eGO term for the gene set are shown under the gene numbers. *P*-values for the eGO analysis are shown in the color bar (top). Full eGO information is listed in Supplementary Dataset 3. **(B)** Heat map using Z-score for expression of all of the genes identified in **(A)**.

We next analyzed the enriched gene ontology (eGO) to determine the biological processes in which each gene set is related (Figure 3A). For instance, there were 139 genes whose expression was down-regulated at 3 h compared to 0 h (q < 10^−10^, Figure 3A and Supplementary Dataset 2). ‘Response to stress’ was the most enriched category for the 139 genes (Figure 3A, Fisher’s exact test p < 10^−9^, Supplementary Dataset 3). ‘Defense response’, ‘response to stimulus’, and other biological processes related to ‘response to stress’ were also enriched categories among the 139 genes (Supplementary Dataset 3). The 211 up-regulated genes from 0 h to 3 h (q < 10^−10^) significantly contains genes categorized as ‘response to chemical’ (p < 10^−28^), ‘response to stimulus’ (p < 10^−24^), and other biological processes (Figure 3A and Supplementary Dataset 3). The eGO analysis suggested that genes in the ‘response to stress’ and ‘response to chemical’ categories were down-regulated and up-regulated by 3 h after treatment, respectively. Next, by 9 h after treatment, ‘photosynthesis’ and ‘response to chemical’ genes were down-regulated and up-regulated, respectively. Finally, ‘response to stress’ and ‘oxidation-reduction process’ genes were down-regulated and up-regulated by 24 h after treatment, respectively.

Next, we generated a heatmap of the expression profiles of the gene sets to see expression patterns (Figure 3B). There were expression pattern variations. Genes down-regulated by the treatments were roughly grouped into early and mid-late down-regulated gene groups. Genes up-regulated were categorized into early, mid-late, and late up-regulated genes. We did not easily find any genes that were specifically influenced by the nanocarbon molecule treatment in the heatmap. Thus, the expression of these genes was changed in a treatment duration-dependent manner, but not a nanocarbon-dependent one.

### 2.3. Impact of nanocarbon molecules on the transcriptome

Because we had only one biological replicate for each time point due to limited amounts of nanocarbon molecules for treatments, it was not acceptable to identify genes whose expression was statistically changed by nanocarbon treatment. Instead, we compared a nanocarbon-treated sample with the control solvent DMF-treated sample that was harvested at the same time after treatment (e.g., CPP_3h vs DMF_3h). Genes whose expression changed two-fold up or down between the nanocarbon-treated and DMF-treated samples were identified (Figure 4, Supplementary Dataset 4 and Supplementary Figure 2). More than 400 genes were obtained in all comparisons using this strategy (Figure 4A and B). The genes whose expression below 1 CMP were regarded as not abundantly expressed (below 1 CPM, Supplementary Figure 2). By contrast, scatter plots between 0h_Rep.1 and MS_9h showed that mis-expressed genes were also found in over 10 CPM (Supplementary Figure 2). We suspected that our strategy might have contained false-positive genes in the comparison between nanocarbon- and DMF-treated samples, due to the lack of statistics related to sample number; however, we reasoned that if the same biological process, an eGO term, varies in multiple comparisons, then nanocarbon molecules have the potential to influence the eGO terms.

**Figure 4.**
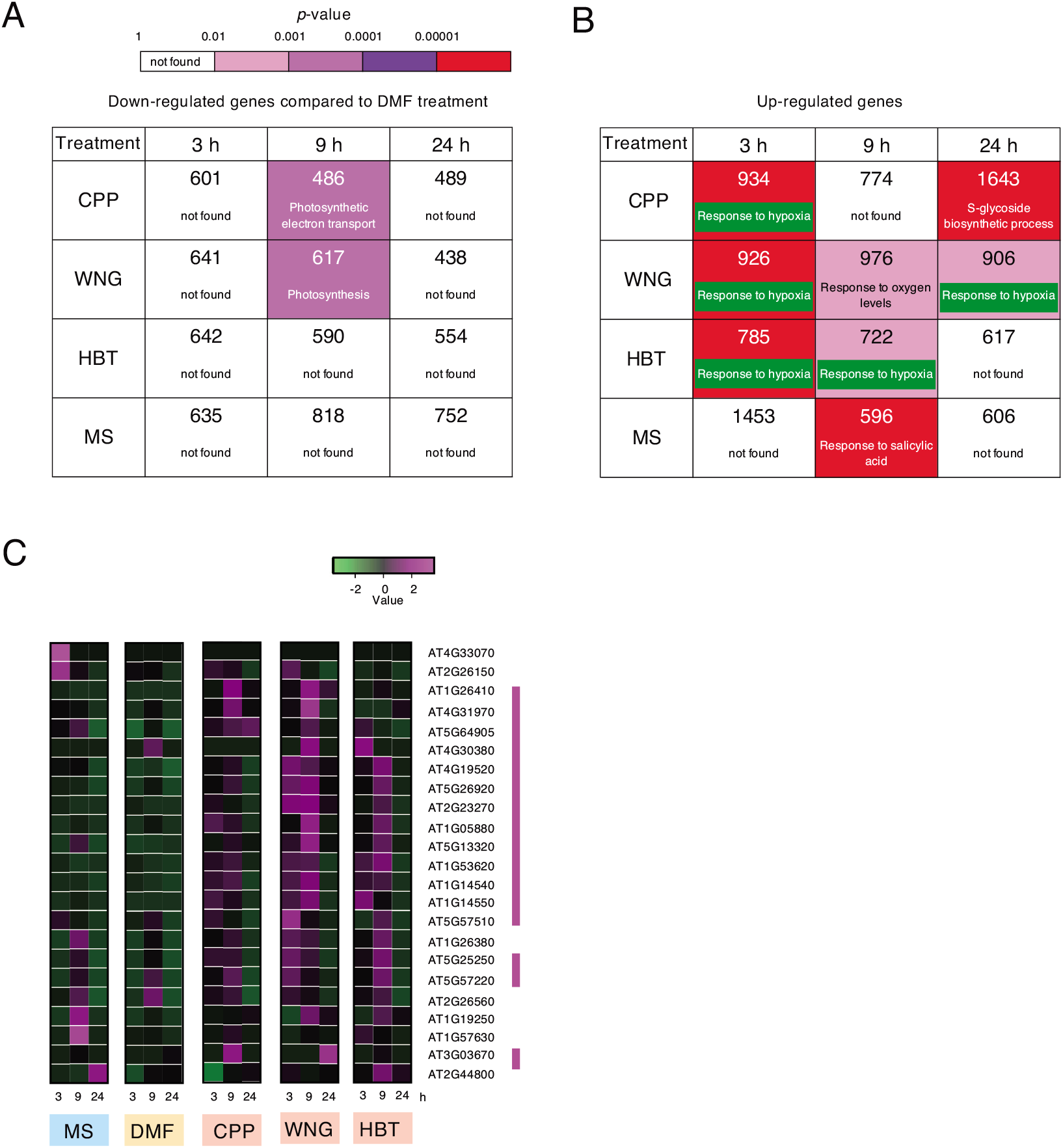
Nanocarbon-dependent alteration of gene expression. **(A)** Number of genes whose expression levels are 2-fold down-regulated by nanocarbon molecules compared with DMF treatment is shown as the upper value in each box. Best hit eGO terms for each gene set appear below the gene number. Full eGO information is listed in Supplementary Dataset 5. **(B)** Number of genes whose expression was 2-fold up-regulated by nanocarbon molecules compared with DMF treatment (upper). eGO for the gene set (lower). The primary gene sets of ‘response to hypoxia’ are indicated by green bars. **(C)** Heat map using Z-score of genes expression annotated to ‘response to hypoxia’. Magenta bars indicate genes upregulated by nanocarbon molecules.

The eGO analysis for ‘over two-fold upregulated genes by 3 h CPP treatment compared to 3 h solvent DMF treatment’ suggested that there is no enrichment in biological processes (p > 0.01) (Figure 4A and Supplementary Dataset 4, Supplementary Dataset 5). Genes downregulated by a 9 h treatment with CPP or WNG significantly enriched genes involved in photosynthesis (p < 10^−3^). Treatment with CPP and WNG for 24 h did not enrich the expression of genes in any biological processes. Genes downregulated by HBT treatment for all time points did not enrich any specific biological processes. The control MS treatment also did not significantly decrease the expression of genes in any specific biological process.

We found that the ‘response to hypoxia (oxygen deficiency)’ gene group was enriched in genes upregulated after a 3 h treatment with CPP, WNG, or HBT, but not water compared to DMF. In addition, a 9 h treatment with HBT caused the upregulation of genes categorized as ‘response to hypoxia’ genes. Treatment of seedlings for 24 h with WNG also upregulated the same gene group. In contrast, treatment of seedlings for 24 h with CPP caused the upregulation of genes related to a different biological process, namely ‘S-glucosinolate biosynthesis’. A 24 h treatment with HBT did not result in significant enrichment of any biological process. Three or 24 h MS treatment compared to DMF treatment did not result in the increased expression of genes involved in any specific biological process. A 9 h treatment with MS resulted in the enrichment of salicylic acid or immune responses. The eGO analysis of different time points after nanocarbon treatment suggested that all three nanocarbon molecules induced a biological response to hypoxia in at least one time point. To test further our hypothesis, we surveyed the expression of genes that are reported to respond to hypoxia (Figure 4C)[25]. The expression of only *AT3G55790* was missing from our analysis. We found that not all genes responding to hypoxia were induced by nanocarbon molecules compared to DMF; however, at least half of the hypoxia responding genes in Arabidopsis were induced by the nanocarbon molecules in RNAseq data.

To validate our finding that nanocarbon-induced genes are among those genes that respond to hypoxia, we conducted a gene expression analysis using RT-qPCR with three biological replicates (Figure 5A). As an internal control, we used expression of *AT3G02780* (*IPP2, ISOPENTENYL PYROPHOSPHATE: DIMETHYLALLYL PYROPHOSPHATE ISOMERASE 2*) gene since *IPP2* expression is constant under many experimental conditions [26]. *IPP2* expression was not changed by nanocarbon molecules in our experiment (Fgure 5B). We found that the expression of *AT5G26920* and *AT2G23270* were not induced by nanocarbon molecules (Figure 5C). Two genes (*AT1G26410* and *AT1G26380*) were induced by nanocarbon molecules, but also by MS treatment, suggesting that the expression of these genes was responding to the treatment. The expression of *AT1G05880 (ARIADNE 12*) was significantly elevated by WNG and HBT treatment compared to the other control experiments (*p* < 0.05). Multiple comparison test did not indicate that CPP treatment induce *ARIADNE 12* compared to control experiments (*p* > 0.05). Thus, *ARIADNE 12* may be usable as a marker for treatment of WNG and HBT.

**Figure 5.**
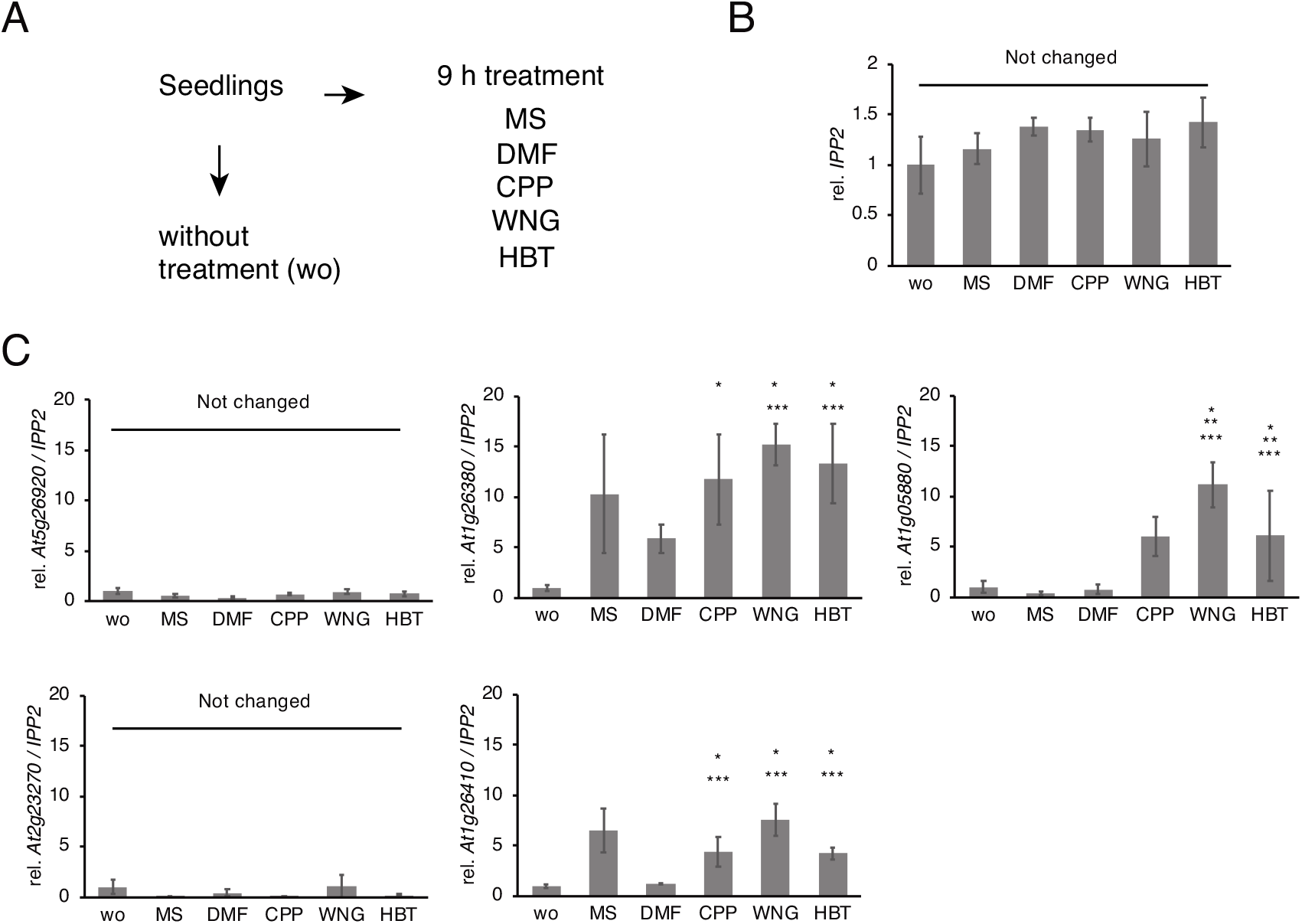
Expression analysis of ‘response to hypoxia’ genes by RT-qPCR. **(A)** Sampling scheme for gene expression analyses. Three independent biological replicates were used. **(B)** Expression of *IPP2* gene used as an internal control. **(C)** Expression of ‘response to hypoxia’ genes. Single, double, and triple asterisks indicate significant difference compared to ‘wo’, ‘MS’, or ‘DMF’ controls (*p* < 0.05, by Williams’s test). Values of ‘wo’ sample were set to 1.0 (**B** and **C**).

## 3. Discussion

Treatment of Arabidopsis seedlings with the nanocarbon molecules did not have an effect on the transcriptome that was as significant as we predicted. Indeed, clustering analysis using all of the transcriptome data indicated that the effect of ‘duration’ had a larger impact than that of ‘nanocarbon molecules’ (Figure 2). This result seems to indicate that using the nanocarbon molecules to control plant physiology or create genetic modifications seems safe from the point of view of toxicity. Cell viability tests using fluorescein diacetate supported the finding that 7-8 h nanocarbon treatments did not result in severe damage to Arabidopsis seedlings (Supplementary Figure 3).

We also conclude that *ARIADNE 12* gene whose transcript was up-regulated is a kind of biomarker for WNG and HBT exposure. *ARIADNE 12* is categorized in hypoxia-induced gene group, and hypoxia is induced when plants are challenged by low O_2[25]_. We hypothesize two possibilities to explain why *ARIADNE 12* is induced by WNG and HBT. First, WNG and HBT localized and accumulated in the apoplastic regions due to their hydrophobic properties, causing the entrapment of O2 there. Secondly, nanocarbon molecules may have attenuated cellular metabolism due to their properties, resulting in a lower amount of O2. Larger nanocarbon molecules (WNG) were more effective than the smaller one (HBT) in influencing the transcription of *ARIADNE 12*. It is still largely unknown how nanocarbon molecules enter plant cells, but we speculate that nanocarbon molecules affecting cellular metabolism must enter cells via the endocytosis pathway.

Plants are frequently challenged by chemical compounds produced by various organisms. Plants recognize such molecules as an indication of invaders and trigger defense responses. Major responses include the salicylic acid- or jasmonic acid-dependent pathways[27]. Given that these pathways were not activated by nanocarbon treatments suggests that Arabidopsis did not recognize the nanocarbon molecules as invaders. Nanocarbon structures are not similar to the structures of molecules found in nature, so plants seem to have no system for responding to these molecules. Our transcriptome analyses also showed no sign of apoptosis or necrosis resulting from nanocarbon treatment. These results highlight the potential for using nanocarbon molecules as nanomaterials to control plant activities in the future.

## 4. Materials and Methods

### 4.1. Preparation of nanocarbon molecules

CPP[28], WNG[29], and HBT[15] were synthesized by our previously reported methods and stocked in our curated nanocarbon library. Each nanocarbon molecule was dissolved in *N,N*-dimethylformamide (DMF, 13015-75, Nacalai) at concentration of 1.0 mM and stocked as a solution at −30°C.

### 4.2. Treatment of Arabidopsis with nanocarbon molecules

Seeds of Arabidopsis thaliana Columbia-0 (Col-0) accession were sown on half-strength Murashige-Skoog (MS) plates[30] (392-00591, Fujifilm-Wako) containing 0.25% (w/v) sucrose (196-00015, Fujifilm-Wako) and 0.3% (w/v) gellan gum (075-03075, Fujifilm-Wako), as previously reported[16]. Plates were kept at 4°C for two days. Four-day-old seedlings grown under constant white light (70 μmol s^−1^ m^−2^) were transferred to 96-well plates (136101 Nunc MicroWell White plate, ThermoFisher Scientific) with a 6 mm diameter dropper. Twenty μL of full-strength MS liquid with 2% (w/v) sucrose and 50 μM nanocarbon molecule dissolved in 5% (v/v) (final concentration) DMF were dropped on the seedlings. Seedlings treated without DMF were prepared as MS-treated samples. Note that all organs were exposure to the treatment. After 3, 9, or 24 h of treatment, 8 seedlings were pooled as a single sample in a 2.0 mL tube containing zirconia beads and frozen with liquid nitrogen. Three biological replicates before treatment were also frozen as time 0 samples. Samples were stored in a −80°C freezer until use. The frozen seedlings were then crushed with zirconia beads, and total RNA was purified. The quality of RNA samples was accessed by measuring the absorbance of OD260 and OD280. The ratio of OD260/OD280 of each sample was higher than 1.9, proving the high quality of RNA sample. Libraries of RNA samples for next generation DNA sequencing (NGS) were generated by previously reported method[16,31].

### 4.3. RNAseq analysis

High throughput DNA sequencing of the library was conducted by Macrogen Japan Corporation with an Illumina platform. The length of each read was 51 bp (single-end). Total reads, Q20 and Q30 values are shown in Supplementary Table 1. Q20 and Q30 indicate 99% and 99.9% accuracy of base calling during sequencing. Resulting reads were obtained in the FASTQ format file. RNAseq data were deposited in DDBJ (http://www.ddbj.nig.ac.jp/) under BioProject ID PRJDB8887.

### 4.4. RNAseq data analysis

The FASTQ file was mapped to Arabidopsis TAIR10 transcripts by Bowtie[32], as previously reported[31]. Mapping rates were over 95% (Supplemental Table 1). Mapped reads were normalized as counts per million (CPM). Clustering analysis of the dataset was performed by pvclust in R[33]. EdgeR in Degust[34] was used to find genes whose expression levels were differently expressed as a function of treatment duration. Enriched gene ontology analyses were performed in PANTHER[35]. Heatmaps were generated by genefilter[36] and gplots[37] in R, as previously described[38,39].

### 4.5. RT-qPCR analysis

Treatment of Arabidopsis seedlings with nanocarbon molecules and RNA extraction were conducted as same for preparation for RNAseq samples. Reverse transcription and qPCR were performed as previously described[16,31]. Amplicon was quantified by standard curve method. *AT3G02780* (*IPP2, ISOPENTENYL PYROPHOSPHATE: DIMETHYLALLYL PYROPHOSPHATE ISOMERASE 2*) gene was used to as an internal control due to its constant expression under many conditions [26]. Expression of other genes were divided by that of *IPP2.* The expression value was further normalized in that of ‘wo’ sample as set to 1.0. Multiple comparison test (Williams’ test) between nanocarbon molecule treatment and control experiments (wo, MS, or DMF) was conducted by detecting significant difference of 0.05. Primers for qPCR are listed in Supplementary Table 2.

### 4.6. Fluorescein diacetate (FDA) staining analysis

FDA treatment to assess the viability of guard cells was examined as reported in a previous study[40] with minor modifications. Arabidopsis seedlings were treated with nanocarbon molecules or 70% (v/v) ethanol for 7-8 h, as described before. Seedlings were then vacuum-infiltrated with 1 μg/ml FDA (069-05451, Fujifilm-Wako) solution in a 10 ml plastic syringe. After 7 min incubation at room temperature, the FDA solution was washed out and the fluorescence emission of guard cells in the abaxial epidermis of cotyledons was detected using a fluorescence microscope (BX-50, Olympus).

## Supplementary Materials

The following are available online at www.mdpi.com/xxx/s1, Figure S1: Circadian clock genes expression, Figure S2: Scatter plots of RNAseq data, Figure S3; The viability of Arabidopsis seedlings after nanocarbon treatment, Table S1: Total reads and Q20/Q30 values of RNAseq data, Table S2; Primers for qPCR analysis, Dataset S1: RNAseq data of plants treated with nanocarbons, Dataset S2; Genes whose expression was significantly (FDR < 1E-010) changed in a treatment-duration manner are listed. Dataset S3; Gene ontology enrichment analysis for genes whose expression are altered a treatment duration-dependent manner, Dataset S4; Gene ontology enrichment analysis for genes whose expression are altered a treatment duration-dependent manner, Dataset S5; Gene ontology enrichment analysis for genes whose expression are altered by nanocarbon treatment.

## Author Contributions

Conceptualization, N.N. A.S., and K.I.; methodology, N.N., A.S., and Y.A.; software, N.N.; validation, N.N., A.S., and Y.A.; formal analysis, N.N. and Y.A.; investigation, N.N. and Y.A.; resources, A.S., Y.S., K.A., and K.I.; data curation, N.N. and Y.A.; writing–original draft preparation, N.N. and A.S.; writing-review and editing, N.N., A.S., and K.I.; visualization, N.N. and Y.A; supervision, N.N., A.S., T.K., and K.I.; project administration, N.N., A.S., T.K., Y.S., and K.I; funding acquisition, N.N., T.K., Y.S., and K.I. All authors have read and agreed to the published version of the manuscript.

## Acknowledgments

We thank Ms. Hiromi Matsuo for part of RT-qPCR analysis, Drs. Masayoshi Nakamura and Naoyuki Uchida for discussion about application of nanocarbon molecules for plants.

