## Supplementary material for "Effect of nanocarbon molecules on the *Arabidopsis thaliana* transcriptome": Supplementary information.pdf

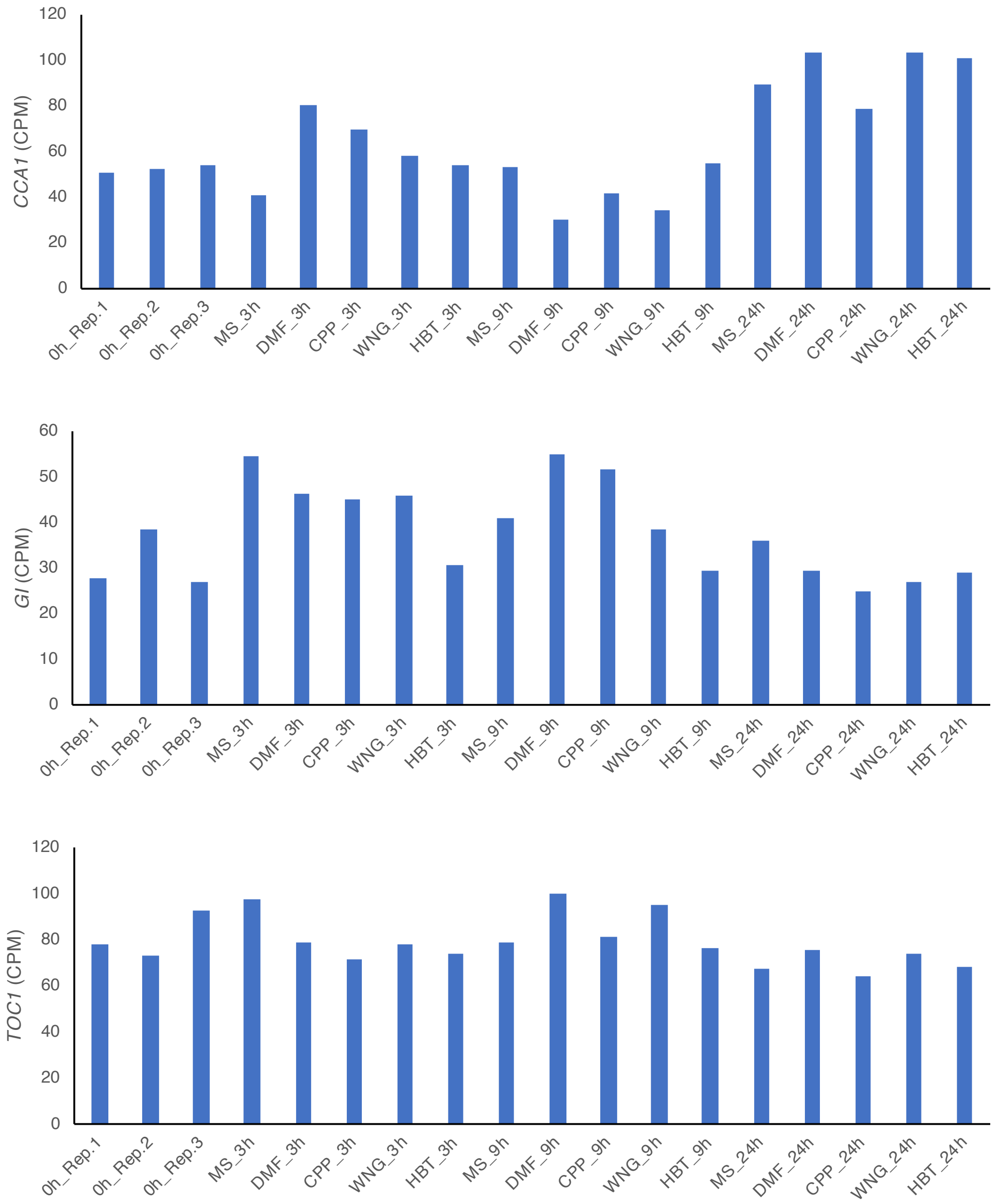

**Supplementary Figure 1. Circadian clock genes expression.** Count per million (CPM) of *GI*, *TOC1*, and *CCA1* in seedlings treated with nanocarbon were shown.

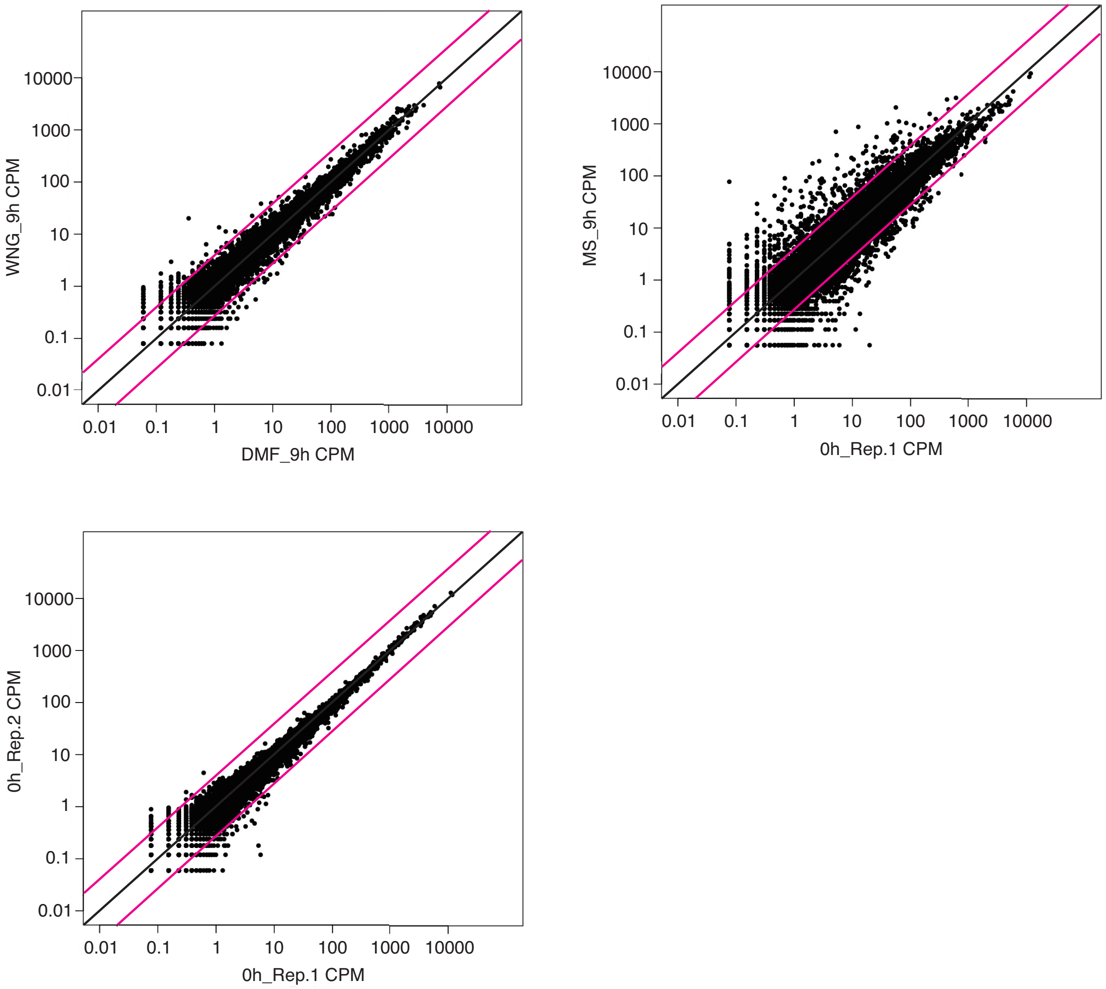

**Supplementary Figure 2. Scatter plots of RNAseq data.**
Transcript levels outside the red lines increased by 2-fold. The comparison between samples treated with MS for 9 h (MS_9h) and the untreated control (0h_Rep.1) had many genes whose transcript levels differed.

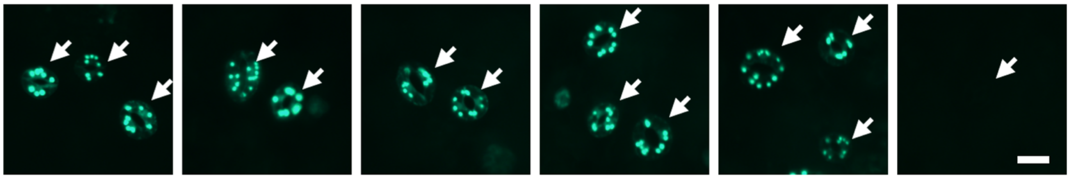

**Supplementary Figure 3. The viability of Arabidopsis seedlings after nanocarbon treatment.**
Seedlings were treated with nanocarbons or 70% ethanol (EtOH) for 7-8 h and then treated with fluorescein diacetate (FDA). Arrows indicate guard cells in the abaxial epidermis of cotyledons. Fluorescence of the guard cells reflects their esterase activity that hydrolyzes FDA to yield fluorescein. Seventy-percent ethanol was used as a control to kill the cells. Representative images from an experiment replicated twice with different biological samples are shown. Scale bar = 20 $\mu$m.

**Supplemental Table1. Total reads, GC/AT% Q20 and Q30.**

| Sample | Total read bases (bp) | Total reads | GC(%) | AT(%) | Q20(%) | Q30(%) | mapping rate (%) |
|---|---|---|---|---|---|---|---|
| 0h_Rep.1 | 685,821,837 | 13,447,487 | 46.7 | 53.3 | 97.27 | 94.29 | 97.2 |
| 0h_Rep.2 | 893,470,683 | 17,519,033 | 46.91 | 53.09 | 97.62 | 94.95 | 95.7 |
| 0h_Rep.3 | 857,941,074 | 16,822,374 | 46.76 | 53.24 | 97.38 | 94.5 | 96.4 |
| MS_3h | 801,485,043 | 15,715,393 | 46.28 | 53.72 | 97.5 | 94.62 | 97 |
| MS_9h | 936,024,828 | 18,353,428 | 46.65 | 53.35 | 97.43 | 94.56 | 96.8 |
| MS_24h | 875,410,665 | 17,164,915 | 46.38 | 53.62 | 97.52 | 94.67 | 97.4 |
| DMF_3h | 553,114,788 | 10,845,388 | 46.62 | 53.38 | 97.39 | 94.54 | 95.5 |
| DMF_9h | 881,942,337 | 17,292,987 | 46.43 | 53.57 | 97.51 | 94.64 | 96.7 |
| DMF_24h | 845,095,857 | 16,570,507 | 46.28 | 53.72 | 97.76 | 95.15 | 96.3 |
| CPP_3h | 791,979,765 | 15,529,015 | 46.82 | 53.18 | 97.41 | 94.41 | 96.9 |
| CPP_9h | 694,792,941 | 13,623,391 | 46.44 | 53.56 | 97.37 | 94.45 | 96.4 |
| CPP_24h | 831,887,112 | 16,311,512 | 46.34 | 53.66 | 97.36 | 94.45 | 96.9 |
| WNG_3h | 625,728,231 | 12,269,181 | 46.45 | 53.55 | 97.33 | 94.42 | 96.5 |
| WNG_9h | 665,656,335 | 13,052,085 | 46.34 | 53.66 | 97.29 | 94.43 | 96 |
| WNG_24h | 609,224,070 | 11,945,570 | 46.19 | 53.81 | 97.39 | 94.58 | 96.1 |
| HBT_3h | 692,107,026 | 13,570,726 | 46.8 | 53.2 | 97.28 | 94.46 | 93.8 |
| HBT_9h | 700,321,749 | 13,731,799 | 46.19 | 53.81 | 97.29 | 94.36 | 96.7 |
| HBT_24h | 513,273,078 | 10,064,178 | 46.21 | 53.79 | 97.25 | 94.25 | 97 |

**Supplemental Table 2. Primers for qPCR analysis.**

| Name | Sequece |
|---|---|
| At2g23270sygF | 5'-GTTCTTCGATGGTTTGTCACTTG-3' |
| At2g23270sygR | 5'-CATTAACTGGTTCGGTCGGG-3' |
| At1g05880sygF | 5'-GCTGAAGCCACACTTCTGC-3' |
| At1g05880sygR | 5'-CTTATGTGGGTGTGATTCGCC-3' |
| At1g26380sygF | 5'-GTTAAAGCTAAGAGTGATCCTGAG-3' |
| At1g26380sygR | 5'-CCAATATTTCAACCTTTTATCTATGC-3' |
| At1g26410sygF | 5'-GGAAGCAATCCAAGTGGTGAG-3' |
| At1g26410sygR | 5'-GCCTATAACACATGCCCAGATAAC-3' |
| At5g26920sygF | 5'-GTTTTCATGGGGGTAGTGGTTAC-3' |
| At5g26920sygR | 5'-GACTACGCTGTATCTCCTCTCGG-3' |
